# A cell type-specific expression signature predicts haploinsufficient autism-susceptibility genes

**DOI:** 10.1101/058826

**Authors:** Chaolin Zhang, Yufeng Shen

**Affiliations:** Department of Systems Biology, Columbia University, New York NY 10032, USA; Department of Biochemistry and Molecular Biophysics, Columbia University, New York NY 10032, USA; Center for Motor Neuron Biology and Disease, Columbia University, New York NY 10032, USA; Department of Biomedical Informatics, Columbia University, New York NY 10032, USA; JP Sulzberger Genome Center, Columbia University, New York NY 10032, USA

**Keywords:** autism spectrum disorders, autism-susceptibility genes, *de novo* mutations, cell type-specific expression signature, D score

## Abstract

Recent studies have identified many genes with rare *de novo* mutations in autism, but a limited number of these have been conclusively established as disease-susceptibility genes due to lack of recurrence and confounding background mutations. Such extreme genetic heterogeneity severely limits recurrence-based statistical power even in studies with a large sample size. In addition, the cellular contexts in which these genomic lesions confer disease risks remain poorly understood. Here we investigate the use of cell-type specific expression profiles to differentiate mutations in autism patients or unaffected siblings. Using 24 distinct cell types isolated from the mouse central nervous system, we identified an expression signature shared by genes with likely gene disrupting (LGD) mutations detected by exome-sequencing in autism cases. The signature reflects haploinsufficiency of risk genes enriched in transcriptional and post-transcriptional regulators, with the strongest positive associations with specific types of neurons in different brain regions, including cortical neurons, cerebellar granule cells, and striatal medium spiny neurons. Based on this signature, we assigned a D score to all human genes to prioritize candidate autism-susceptibility genes. When applied to genes with only a single LGD mutation in cases, the D score achieved a precision of 40% as compared to the 15% baseline with a minimal loss in sensitivity. Further improvement was made by combining D score and mutation intolerance metrics from ExAC which were derived from orthogonal data sources. The ensemble model achieved precision of 60% and predicted 117 high-priority candidates. These prioritized lists can facilitate identification of additional autism-susceptibility genes.

## Introduction

Autism or autism spectrum disorders (ASDs) are very common neurodevelopmental diseases characterized by deficits in language, impaired social interaction, and repetitive behaviors with complexes such as seizures and intellectual disability (1, 2). Symptom onset is typically early (~3 years old) and the current estimate of incidence is over 1% worldwide (3), making ASDs a huge burden for affected families and for society.

Genetic risk factors are believed to play a pivotal role in ASDs, as revealed by a concordance rate up to 90% between monozygotic twins and by over 10-fold increase in the risk for a new born child if a previous sibling is affected (4).Some syndromic forms of autisms are known to be monogenic, as represented by mutations of *FMR1* (encoding FMRP) in the fragile X syndrome that is comorbid with autism and accounts for up to 5% of ASD cases (5, 6). However, most of the genomic abnormalities or mutations found in autism patents are extremely rare and frequently *de novo*. Earlier studies using microarray-based approaches identified hundreds of *de novo* copy number changes (CNVs) (7–12). More recently, *de novo* mutations in individual nucleotides, including single nucleotide variations (SNVs) and small insertions and deletions (indels) were identified by exome-sequencing (13–18) or whole genome sequencing (19, 20).

The exciting progress in these genetic studies has provided important insights into the etiopathogenesis of autism. First, at least a substantial proportion of autism risk is conferred by individually rare mutations affecting one or more disease-susceptibility genes. The number of risk loci has been estimated to be in the range of several hundred to over 1,000 genes (4, 16, 17). Second, although the complexity of the genetic landscape underlying autism is still a matter of debates, one theory supported by several lines of evidence favors that a large number of autism risk loci are individually of high penetrance (4, 21, 22). Third, analysis of the seemingly isolated candidate autism-susceptibility genes points to disruption in several convergent molecular pathways (4, 16, 23–25) that inform the neurobiological underpinnings of autism (reviewed by ref. (26)).

Strikingly, even the most frequent *de novo* mutations in single genes can explain no more than 1% of ASD cases (26). This extreme genetic heterogeneity presents a big challenge for conclusive identification of autism-susceptibility genes, which impeded further functional studies of autism neurobiology and development of therapeutic strategies. A particular group of *de novo* mutations identified by whole exome-sequencing (13–16, 18) are “likely gene-disrupting” or LGD mutations, which is a collection of severe mutations introducing frame shift, disrupted splice sites or premature stop codons. Probands clearly show a higher burden of *de novo* LGD mutations than their unaffected siblings used as control, indicating enrichment of disease-susceptibility genes disrupted by these genomic lesions. However, although about 4,000 ASD patients and their families have been sequenced so far, only about 40 genes at best have been determined as high-confidence autism-susceptibility genes based on their recurrence. Most of the remaining mutations were observed in only single patient, and ~80% of these are expected to be non-pathogenic (see Material and Methods). The signal-to-noise is even lower for genes with missense mutations (17) due to their more moderate effects and high background mutation frequency. Therefore, most autism-susceptibility genes are currently buried among a growing list of potential candidates.

Two general strategies can be used to facilitate the identification of candidate autism-susceptibility genes. One strategy is to sequence larger cohorts of ASD families. For example, the SPARK project (27) aims to recruit and analyze 50,000 individuals of autism and their families. Furthermore, whole-exome or whole-genome sequencing is also complemented by targeted re-sequencing to reduce cost (28). A caveat of this strategy is its prohibitive cost associated with recruitment and sequencing of large cohorts. A second complementary strategy is to stratify already identified mutations based on existing orthogonal information associated with the affected genes. For example, depletion of rare deleterious variants estimated from the general population reflecting severe selection pressure is effective in prioritizing deleterious mutations (29–31). Alternatively, gene regulatory and function information can also be used to help distinguish pathogenic vs. neutral mutations. The latter is based on the assumption that the common clinical phenotype of ASDs originates from certain common features shared by autism risk loci at the molecular level. Along this line, recent studies identified molecular pathways underlying autism etiopathology from analysis of shared genetic phenotypes (24, 32), protein-protein interactions (14), and gene co-expression networks (23, 25). However, these previous studies utilizing network analysis techniques were not designed to optimize the prioritization of individual autism-susceptibility genes *per se*, or to rigorously evaluate the signal-to-noise ratio associated with prediction.

In this work, we predict autism-susceptibility genes by using gene expression profiles in a wide range of specific neuronal cell types, with the assumption that different cell types have different susceptibility or relevance to autism etiopathogy. Due to the heterogeneity among many different cell types in the brain, such an analysis may reveal cell type-specific gene regulation that cannot be detected by analysis of brain tissues used in many previous studies. We identified a gene expression signature reflecting haploinsufficiency in the context of autism that was able to effectively predict whether individual LGD mutations confer disease risk. Importantly, the use of cell type-specific expression also allows us to highlight the cellular contexts of the identified risks. Furthermore, this signature is complementary to previous mutation intolerance analysis, and an ensemble model combining multiple scoring metrics gave the optimal prediction accuracy.

## Results

### DAMAGES analysis uncovers an expression signature of autism-susceptibility genes

Our study was motivated by a postulation that different cell types in the brain have different susceptibility and impact on autism etiopathology,which is supported by recent studies showing expression bias in candidate autism-susceptibility genes (32, 33). However, to the best of our knowledge, the effectiveness of cell type-specific expression in predicting autism-susceptibility genes, alone or in combination with other metrics, has not been systematically explored.

We developed and applied a computational framework for disease-associated mutation analysis using gene expression signatures (DAMAGES) to score the association of human genes with autism, to refine the lists of candidate autism-associated genes currently available, and to uncover features shared by these genes as a means of understanding the molecular underpinnings of the disease (Figure S1). In contrast to previous network analysis approaches, we adopted a case-control classification framework so that it can take advantage of rich resources of existing gene expression profiles and machine learning approaches to optimize and objectively evaluate the accuracy of prediction. To provide a proof of principle in this study, we focused on the utilization of gene expression profiles in a wide range of isolated central nervous system (CNS) cell types. We decided to examine a large microarray dataset that profiles cell type-specific transcripts associated with translating ribosomes in the mouse brain generated by a biochemical assay named TRAP (34, 35). In total, this dataset is composed of translational profiles of 24 specific mouse CNS cell types, including both neurons and glial cells, isolated from six different regions, together with unselected RNA representing all cell types in each of these regions (34) (Figure S1 A,C). This translational profiling approach was previously shown to give robust gene expression measurements, and to effectively identify known and novel cell-type specific markers and provide biological insights into each cell type (34).

To identify expression signatures of autism-susceptibility genes, we started with a list of 162 genes containing *de novo* LGD mutations in either ASD probands or unaffected siblings collected from four exome-sequencing studies (13–16) (Figure S1B). Our prediction model was built using these mutations representing all information available before 2013, which gave us a chance to use additional genes discovered by large-scale followup studies for objective evaluation. In total, 145 genes have mouse orthologs and were represented in the microarray dataset we used (Table S1). We assume that the 33 genes with LGD mutations in unaffected siblings confer no risk of ASD (non-disease genes). On the other hand, the other 112 genes with LGD mutations in probands represent a mixture of ~65 disease-susceptibility genes and ~47 non-disease genes (Table S1 and Material and Methods; a similar estimate provided in (25)). It is worth noting that we limited our analysis to mutations derived from unbiased genomic screens to build the model, and excluded candidates identified by more targeted or hypothesis-driven approaches or by transcriptomic analysis to avoid potential ascertainment bias. This is particularly critical for an objective assessment of DAMAGES analysis in prediction accuracy.

Given the relatively balanced representation of autism-susceptibility genes and non-disease genes (estimated to be ~65 and ~80, respectively) in the dataset, we anticipated that the contrast between these two groups of genes would represent a major axis of expression dynamics in the high-dimensional space. A principal component analysis (PCA) (36) was thus performed to identify the orthogonal axes that explain the most variance. This analysis revealed that projection of genes to the second principal component (PC) is very predictive of mutations in probands versus controls (Figure 1A-B). We note that PCA is an unsupervised method which does not incorporate information on the source of mutations so the prediction performance is not due to data overfitting. To have a more rigorous assessment of the ability of each PC or combination of PCs to differentiate potentially disease-associated versus neutral mutations, we performed a regularized linear regression analysis lasso (37) to find the PCs that are most predictive of the source of mutations. In this method, a parameter λ, specifying the penalty against more complex models, controls the number of PCs contributing to the regression model (i.e., having a non-zero regression coefficient β). We found that only PC2 has a non-zero coefficient across a wide range of λ, while the coefficients of all other PCs shrink to zero very quickly as λ increases (Figure 1C). To determine whether the regression models overfit, we performed leave-one-out cross validation (LOOCV) to predict the source of mutations using varying values of λ (and thereby different numbers of PCs in the regression models). The best performance was achieved when a single component PC2 was used for prediction (λ=5) (Figure 1D). Therefore, we decided to use the PC2 as a signature of autism-susceptibility genes, and adjusted the threshold according to LOOCV, which resulted in the final DAMAGES scores (or D scores) used for gene ranking. As a result, we were able to identify 93 genes with positive D scores, including 83 genes with LGD mutations in probands and 10 genes with LGD mutations in siblings, respectively (arrowhead in Figure 1B and Table S1). Proband-specific LGD mutations are strikingly enriched in genes with positive D scores compared to the remaining genes (Figure 1B; odds ratio=6.48, *P*=8.06×10^-6^, Fisher’s exact test). The ability of the gene expression signature to differentiate mutations in probands from those in siblings suggests that at least some of the CNS cell types included in the microarray dataset are strongly associated with the underlying molecular mechanisms of autism.

**Figure 1:**
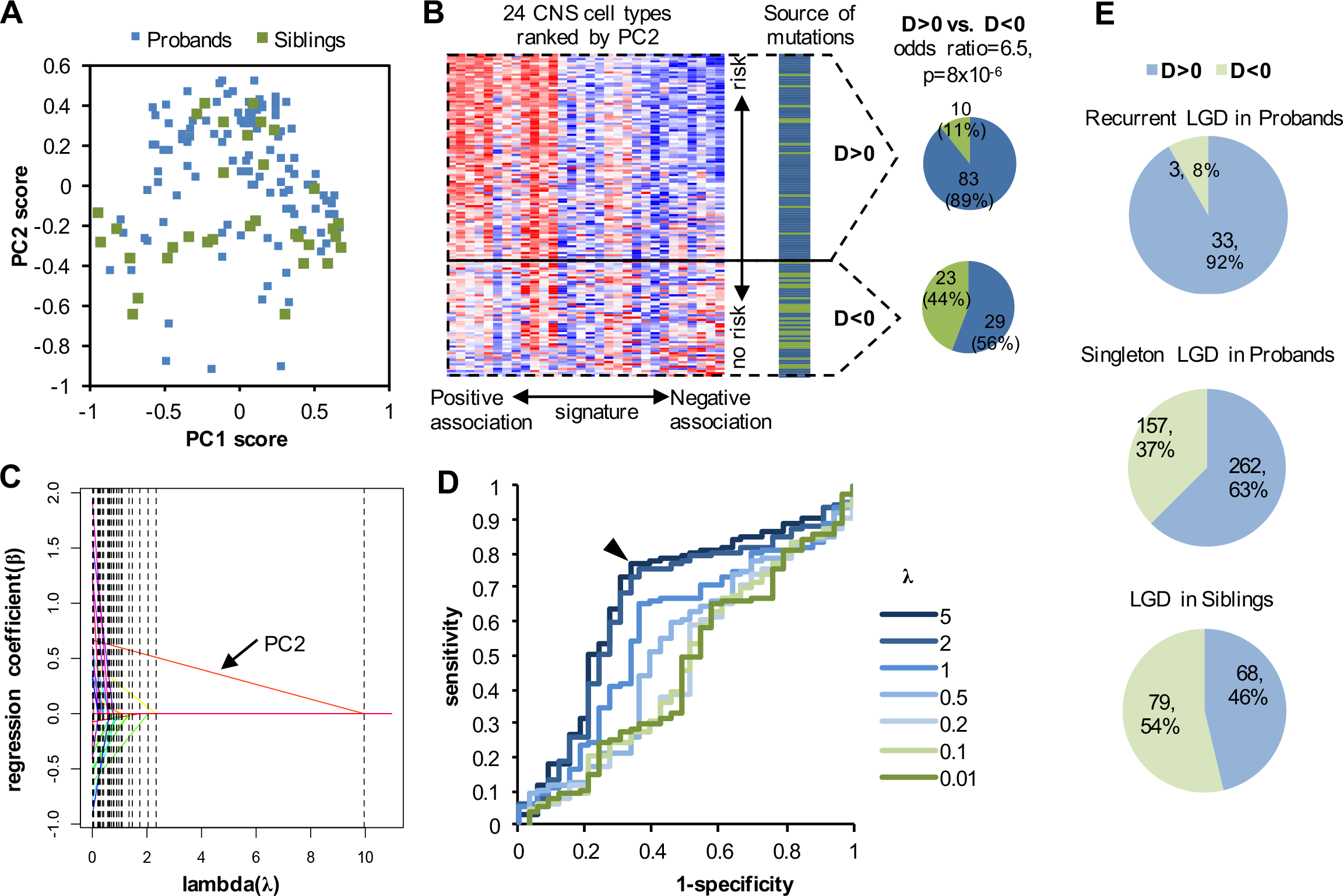
A molecular signature differentiates autism-susceptibility genes and non-disease genes. **A.** A total of 145 genes, including 112 genes with LGD mutations in probands (blue dots) and 33 genes with LGD mutations in siblings (green dots) are projected onto the two-dimensional space defined by the first two principal components (PCs). **B.** The second principal components differentiates autism-susceptibility genes and non-disease genes. In the heatmap on the left, the PC2 score and loading were used to rank genes and arrays respectively. The source of mutation in each gene is indicated with genes shown in the same order. The arrowhead indicates the threshold used for further analysis in this study, and this threshold gives the same list of genes as those predicted by LOOCV using lasso model with λ=5 (panels **C. D** below). The number of genes with D score>0 or <0 is shown on the right. **C.** A regularized regression analysis is used to evaluate the relevance of each PC in predicting the source of mutations (i.e., probands versus siblings). Parameter λ controls the number of non-zero regression coefficients and therefore the complexity of the models. Each curve represents the trace of one coefficient when varying values of λ are used. The trace for the coefficient of PC2 is indicated. Dotted lines indicate shrinkage of coefficients to zero. **D.** Leave-one-out cross-validation (LOOCV) is used to evaluate the specificity and sensitivity in predicting the source of LGD mutations. Prediction is based on genes ranked by the regression score. Each receiving operating characteristic (ROC) curve is derived from one specific value of the regularizing parameter λ. The best performance is achieved when only PC2 is used for prediction (λ=5). The arrowhead indicates the turning point of prediction performance when different thresholds of varying stringency are used for prediction. **E.** Summary of prediction using an expanded list of genes with LGD mutations in ASD patients and unaffected siblings.

### Validation of the expression signature using expanded autism exome sequencing data sets

We assigned a D score to all human genes represented in the microarray dataset independent of their mutation status (Table S2). This allowed us to have an independent, “prospective” evaluation of the performance of the expression signature using an expanded list of genes with LGD mutations from recent large-scale studies after our prediction model was built (Figure 1E and Table S 1)(17, 18). In this expanded dataset, almost all genes with recurrent mutations in autism patients (35/38=92%) received a positive D score (exceptions are *DSCAM*, *RANBP17*, and *TCF7L2*; two genes not represented on the array were excluded). Among the genes ranked in top 25% by the D score, there is a 2.8 fold enrichment (P=3.2×10^-12^) of LGD mutations in cases comparing to unaffected siblings, whereas there is no significant enrichment in rest of genes (rate enrichment = 1.2, P=0.16) (Table 1). Therefore, DAMAGES analysis prioritized *bona fide* autism-susceptibility genes with minimal loss.

**Table 1:**
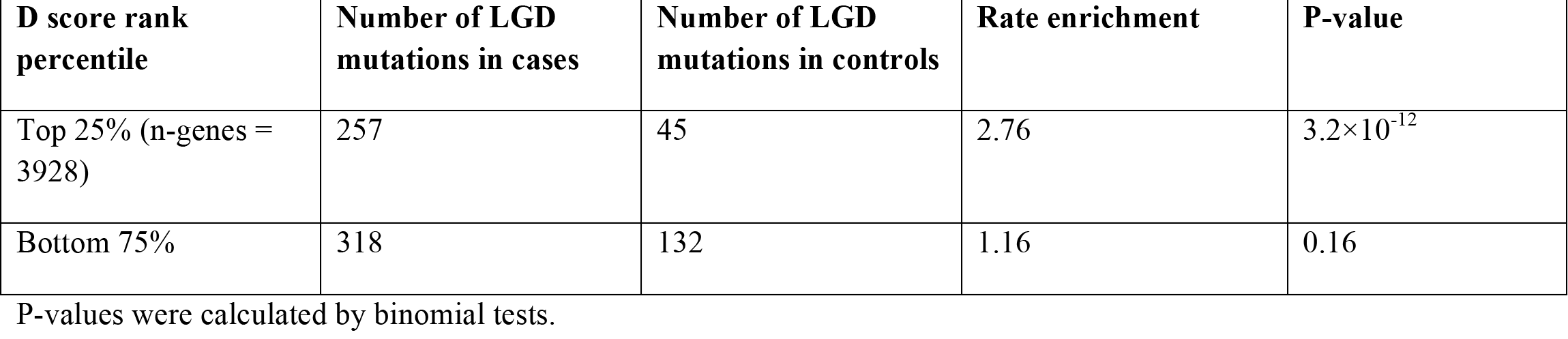
Enrichment of LGD *de novo* mutations in cases among genes grouped by D score.

For additional validation, we examined 528 genes compiled in the Simon Foundation Autism Research Initiative (SFARI) autism gene database, a list of potential autism-associated genes manually curated by experts according to various types of evidence available in the literature (http://gene.sfari.org) (38). Among the 483 SFARI genes represented in the microarray data, 300 (62%) have a positive D score (Figure 2A and Table S3), a very significant enrichment compared to all genes represented in the microarray dataset (odds ratio=2.34, *P*=6.3* × 10^-20^; Fisher’s exact test). In addition, this proportion is higher for genes with more evidence supporting their implication in autism (Figure 2B). For example, genes with positive D scores include 5/5 (100%) genes that are classified as strong candidates and 15/20 (75%) genes that are classified as syndromic, such as *FMR1*,*MECP2* [Rett syndrome (39)] and *TSC1/2* [tuberous sclerosis complex (40)]. Furthermore, SFARI genes received much higher ranks based on the D score, as compared to ranking by the first PC reflecting the neuron-glial distinction (*P*=1.2× 10^-22^; Wilcox ranksum test; see below).

**Figure 2:**
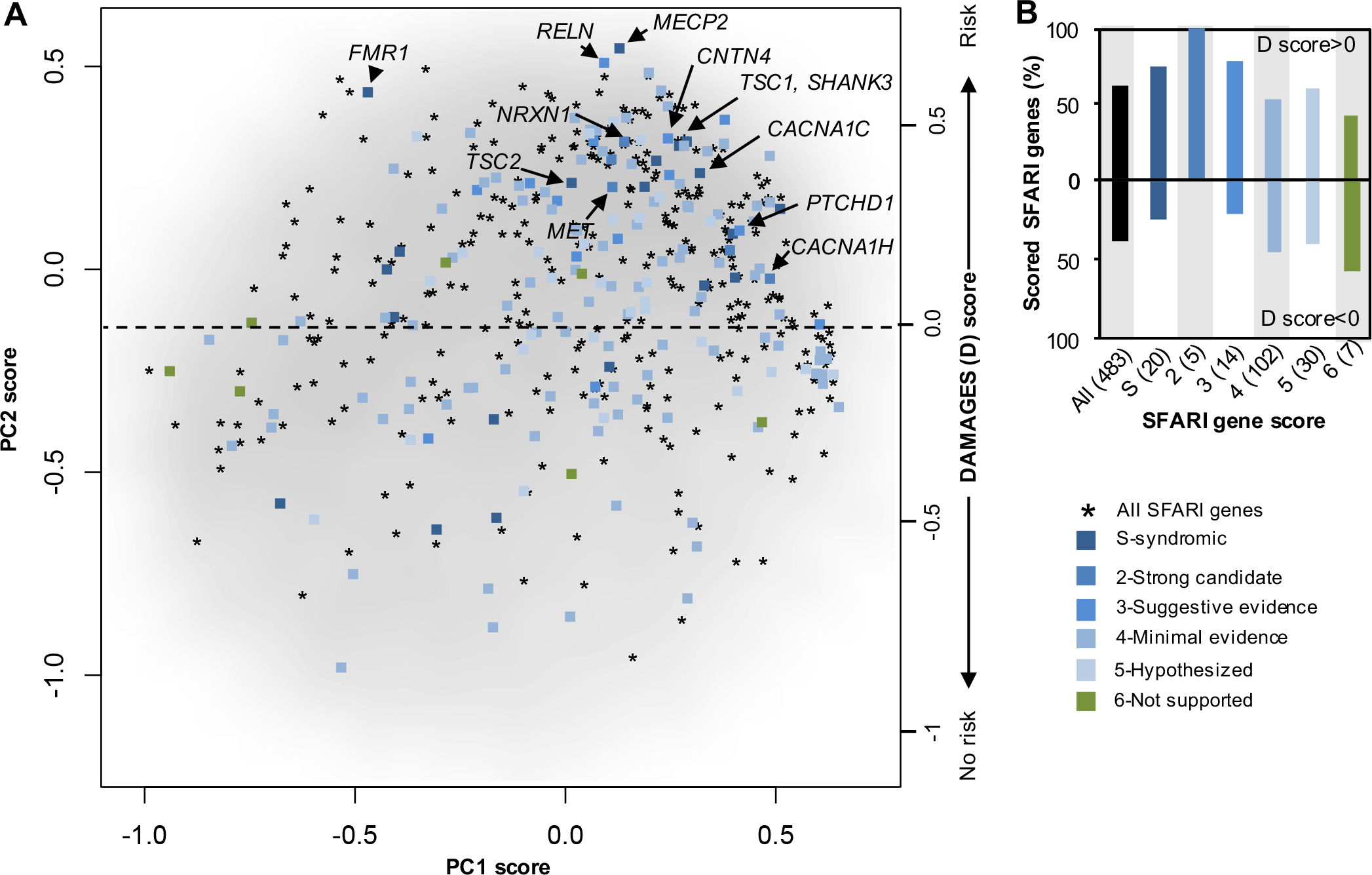
The DAMAGES molecular signature refines the list of candidate genes in the SFARI autism gene database. **A.** All genes represented in the microarray dataset, except those with average log_2_ intensity <4.5, are projected onto the first two PCs, and shown as a smoothed scatter plot. The gray-scale intensity reflects the local density of genes. A total of 483 genes from the SFARI autism gene database represented in the microarray dataset (asterisks) are overlaid. A subset of these genes were manually scored by experts by considering strength of existing evidence, and these scored genes are distinguished using different colors. A subset of syndromic ASD genes and the five strong candidate genes are highlighted. **B.** The percentage of scored genes in each group with a positive or negative DAMAGES score (D score) is shown. The color codes are the same as in (**A**). The number of genes in each group is indicated in the parentheses following the gene categories.

### CNS cell types associated with autism

To further confirm this molecular signature and gain more biological insights, we examined the loadings of different cell types on each PC (Table S4). The first PC essentially differentiates neurons versus glial cells and unselected cell types in different brain regions (Figure 3A). In contrast, the second PC predictive of autism-susceptibility genes appears to give more of a mix of different cell types and regions, although a certain bias is also clear (e.g., cortical neurons have the highest positive loadings; see below). To assess whether this pattern reflects their association with the underlying molecular mechanisms of autism and is specific for autism-susceptibility genes, we performed another PCA using all genes showing the most variation across different cell types. The first PC of the whole dataset is highly correlated with the one derived from genes with LGD mutations (R^2^=0.74), and similarly differentiates neurons from glial cells and unselected cell types (Figure 3B). This result is consistent with the notion that even among genes with LGD mutations the distinction of neuronal versus glial genes dominates the expression dynamics. In contrast, the second PC identified in the global PCA has a low correlation with the second PC identified using genes with LGD mutations (R^2^=0.19; Figure 3C). This observation supports the notion that the molecular signature identified using genes with LGD mutations indeed reflects certain specific features shared by autism-susceptibility genes.

**Figure 3:**
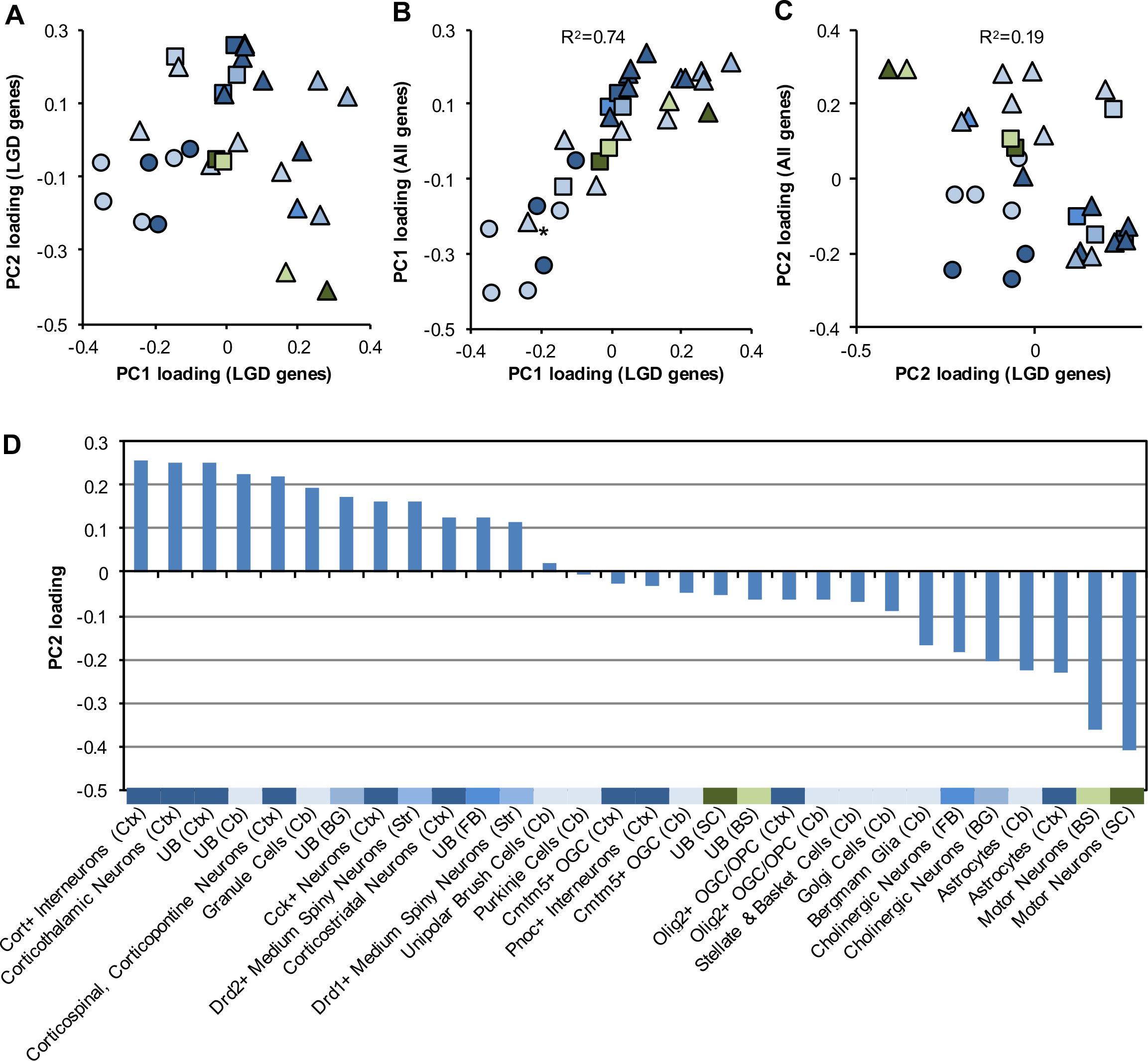
The DAMAGES molecular signature reveals CNS cell types associated with autism. **A.** The loadings of different cell types on the first two PCs derived from genes with LGD mutations are shown. Each dot represents a cell type. Different colors represent the brain regions used to isolate the specific types of cells, with the same color codes as shown in **Figure S1A**. Neurons, glial cells and unselected RNA samples are represented by triangles, circles, and squares, respectively. **B.** The loadings of cell types on PC1 derived from genes with LGD mutations (x-axis) are plotted against that derived from the whole dataset (y-axis). The asterisk indicates cerebellar Grp^+^ cells that are known to include both unipolar brush cells and Bergmann glial cells (34). The squared Pearson correlation between the two signatures is indicated. **C.** Similar to (**B**), except that the loadings on PC2 are plotted. **D.** Loadings of all cell types on PC2 derived from genes with LGD mutations (DAMAGES signature) are plotted. The color codes and abbreviation of each brain region are the same as shown in **Figure S1A**. UB: unbound RNA without selection for specific cell types.

Therefore, the loadings of different cell types on the signature (PC2) likely reflect their association with autism (Figure 3D). In general, none of the glial cell types included for this analysis has a positive association. On the other hand, the association of neurons with the signature varies depending on specific cell types and brain regions. Different types of cortical neurons, including interneurons, corticothalamic neurons, corticospinal and corticopontine neurons, Cck^+^ neurons, and corticostriatal neurons, have large positive loadings on the signature. However, not all types of cortical neurons have a positive association, and some, such as Pnoc^+^ interneurons, have a negative loading. Besides cortical neurons, cerebellar granule cells, striatal medium spiny neurons, but not Purkinje cells, cholinergic neurons, or motor neurons, show a strong positive loading. Altogether, these observations are not only consistent with autism being mainly an impairment of high-level cognitive functions, but also suggest that even in a given brain region, specific cell types may play very different roles in the etiopathogenesis of the disease.

### Molecular functions associated with the autism-susceptibility gene expression signature

We asked whether the expression signature captures certain molecular functions shared by autism-susceptibility genes.To this end, we performed Gene Ontology (GO) analysis (41) using the top 500 protein-coding genes ranked by D scores independent of their mutation status. This analysis revealed very strong enrichment of those involved in “transcription” (Benjamini FDR=3×10^-14^), “chromatin modification” (Benjamini FDR=7.9×10^-6^) and “regulation of RNA metabolic process” (Benjamini FDR=2.2×10^-4^) (Table S5). It is worth noting that “chromatin organization” is also enriched in the 83 high-priority candidate genes, although the statistical significance is marginal (Benjamini FDR=0.07). Therefore, not only are genes with LGD mutations themselves enriched in those important for transcriptional regulation, as noted previously (14, 16, 42), but they define a molecular signature represented by a larger set of genes with coherent molecular functions in both transcriptional and post-transcriptional regulation of gene expression.

### The expression signature reflects haploinsufficiency

We postulate that the expression signature may reflect haploinsufficiency because it was derived from genes with heterozygous loss of function. To test this hypothesis, we examined genes covered by relatively focal *de novo* CNV events (≤50 genes) detected in ASD probands (7–9, 11, 12). Interestingly, genes with higher D scores tend to overlap with deletions than amplifications (P<0.04; Spearman correlation test). The significance is relatively marginal, presumably due to the limited spatial resolution of the CNVs. For further confirmation, we examined genes differentially expressed in post-mortem autism brains as compared to controls (43), assuming that the dosage-dependent alteration can also be caused by changes at the transcription level. Indeed, genes down-regulated in autism tend to have a positive D score (P<2.2×10^-16^), while genes upregulated in autism tend to have a negative D score (P<2×10^-7^; Figure S2A).

Recently the Exome Aggregation Consortium (ExAC) used large-scale exome sequencing data of general populations without developmental disorders to estimate metrics of haploinsufficiency (31), including the probability of being loss-of-function (LoF) intolerant (pLI) and LoF Z-score. A positive correlation (r = 0.29, P < 10^-10^) of LoF Z score and D score was observed among genes with positive D scores (Figure S2B). This observation is again consistent with the notion that both metrics are related to haploinsufficiency although they were derived using entirely different data sources and assumptions.

### Prioritizing genes in CNVs including 16p11.2

*De novo* CNVs detected in ASD probands typically span dozens or hundreds of genes (7–12). Therefore, although over 2,000 genes are covered by at least one CNV identified so far, it is difficult to differentiate *bona fide* autism-susceptibility genes from other passenger genes. We focused on 58 deletion CNVs for which all overlapping genes have mouse orthologs and are represented in the microarray data. Of these, 30 CNVs each have one and only one gene with a positive D score. Based on the high sensitivity (~90%, see below) of a positive D score in predicting autism-susceptibility genes, we argue that if a CNV is pathogenic, the only gene with a positive D score is the most likely causal gene. We therefore denote the CNV “likely supporting CNV” or LS-CNV of the corresponding gene. This analysis resulted in 19 genes supported by deletion LS-CNV events in one or more patients (Table S6). Of these genes, five are supported by recurrent LS-CNVs *(NRXN1, DPP6, PTPRT, SHANK2* and *SLC4A10)*, and all of these five genes are known to have functional implications in synapse (44–48) and/or autism-related phenotypes (47). Remarkably, three genes harbor recurrent LGD mutations in ASD patients (*ANKRD11*, *CHD3* and *KMT2C*; odd ratio=130, P<3.4×10^-6^, Fisher’s exact test). Two additional genes (*NRXN1* and *SHANK2*) have singleton LGD mutations from exome sequencing (i.e., recurrent if the LS-CNV is counted).

LS-CNVs tend to span a smaller number of genes than CNVs in general. For a majority of CNVs overlapping with more genes, it is difficult to reliably distinguish susceptibility genes versus passenger genes even with the D score. Nevertheless, it is still possible to eliminate a substantial fraction of passenger genes. To illustrate this point, we examined the most frequent recurrent *de novo* CNV located in 16p11.2, which accounts for up to 1% of ASD cases (9) (14 deletions and 5 duplications in the dataset used for this analysis; Figure 4A). This region spans 26 genes (11) and all of them have mouse orthologs; deletion of the region in mice phenocopied behavior deficits observed in ASD patients (49). Among the 23 genes represented in the microarray data, nine have a positive D score (Figure 4B). Interestingly, deletion of a smaller region in this locus also segregates with ASD or ASD traits (50). This deletion encompasses five genes, including *KCTD13*, *ASPHD1* and *SEZ6L2* with a positive D score. A recent study further demonstrated that *KCTD13* is a major driver of the macrocephalic phenotype associated with ASD cases carrying the 16p 11.2 CNV (51). Some of the other genes in this locus, especially the ones highlighted by DAMAGES analysis, could contribute to the additional clinical manifestations in ASDs.

**Figure 4:**
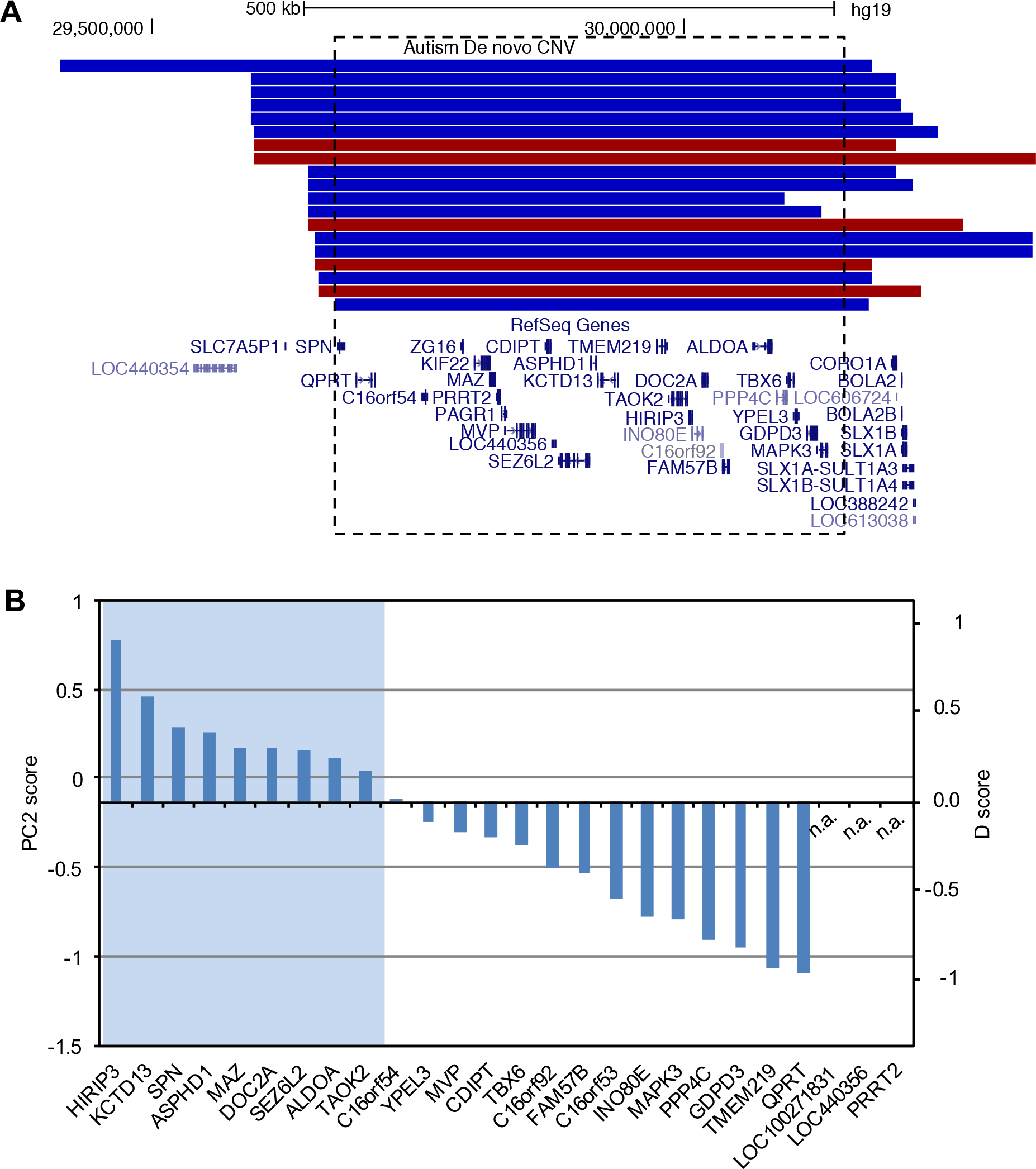
Prioritized candidate autism-susceptibility genes with recurrent CNVs in chromosome 16p11.2. **A.** A UCSC genome browser view of the region (hg19:chr16:29,350,841-30,433,540) is shown, with *de novo* CNV events displayed above the RefSeq genes. Duplication and deletion CNVs are shown in red and blue, respectively. The dotted box indicates the region with 26 genes affected in almost all CNV s. **B.** The 26 genes are ranked by their D scores. Three genes not represented in the microarray data are indicated by n.a.

### An ensemble model for optimized prediction of autism-susceptibility genes

We next compared the performance of D score and ExAC scores in predicting autism-susceptibility genes and investigated if improved performance can be achieved by combining these different metrics. We focused on genes with single LGD mutations in cases and found ExAC pLI>0.9 or LoF Z-score>3 achieved similar optimal performance as D score (about 40% precision and 90% sensitivity; Figure 5A, B), as compared to a baseline 15% precision. Importantly, while predictions by the two metrics overlap, small genes with low background mutation rate (such as MECP2, pLI=0.7) tend to be missed by ExAC scores. To investigate whether ExAC scores and D score make independent contributions in gene prioritization, we performed a logistic regression to classify recurrently mutated genes in cases and genes with LGD mutations in controls, with ExAC pLI, ExAC mis-z, and D score as features. The coefficients of all three features deviate significantly from zero (Table 2), indicating that D score is complementary to ExAC scores in determining gene LGD intolerance (interestingly, gene expression in embryonic mouse brain did not any predictive power). Therefore, combining these scores would maximize the performance in candidate gene prioritization. To assess that, we applied the estimated logistic model to all genes to calculate an ensemble score (Table S2), and estimated precision-recall rates in a range of top rank thresholds, excluding the genes that are recurrently mutated. We found that the ensemble score outperforms all individual methods, with an optimal performance among the top 1,300 genes (Figure 5B,C). With ensemble score, the precision quadruples to 60% with near-maximal sensitivity (estimated to be 97%). Using this threshold, we identified 117 candidate genes with a singleton LGD mutation (Table S7). Since ASD shares substantial number of risk genes with other neurodevelopmental disorders (18, 52), we obtained the *de novo* mutation calls from the latest released data (53) from the Deciphering Developmental Disorders (DDD) project (54) to further assess the performance of ensemble score prediction. The DDD data includes 4,293 patients with severe undiagnosed developmental disorders. Among 117 candidate genes predicted by ensemble score, 65 harbor at least one LGD or damaging missense (predicted by metaSVM (55) or polyphen-2 (56) and CADD (57)) *de novo* mutations in the DDD data set. This rate is much higher than non-candidate genes with a single LGD mutation in ASD data (odds ratio = 4.6; p-value = 2×10^-11^). Conversely, among all the genes with a single LGD mutation in ASD data, the ones with at least one damaging mutation in the DDD data have higher ensemble score than the ones without (P = 1.3×10^-11^; KS test) (Figure 5D). These observations indicate that the candidate genes predicted by the ensemble score are much more likely to be associated with developmental disorders in general.

**Figure 5:**
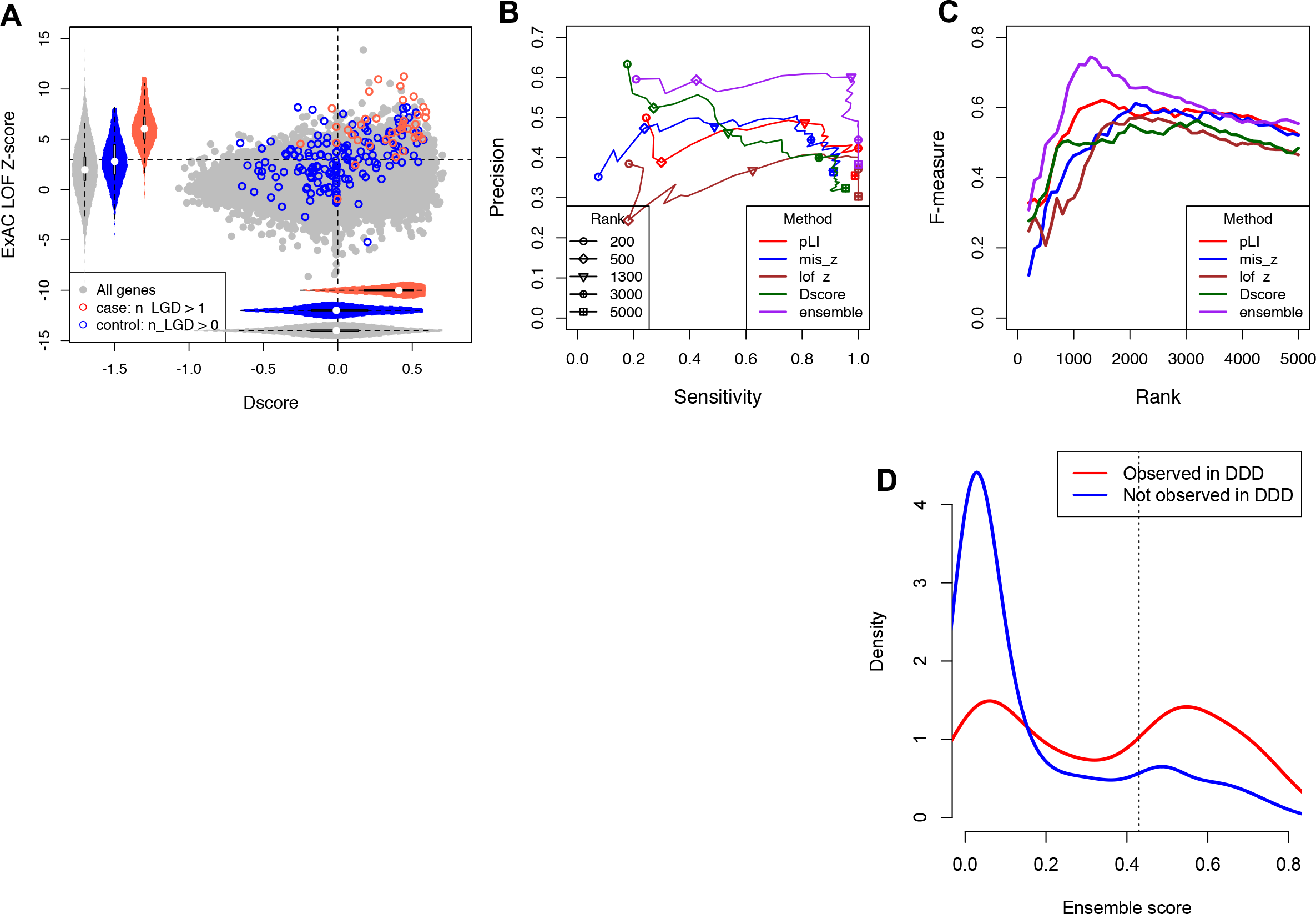
Comparison and integration of D score with ExAC scores. **A.** Distribution of D scores and ExAC LOF Z-scores for genes with recurrent LGD mutations in ASD cases and controls. **B.** Performance of prediction as measured by Precision and Recall using different scoring metrics or an ensemble model as a function of varying cutoffs. **C.** Similar to (**B**), but the F measure is shown. D.Distribution of ensemble score among genes in which damaging *de novo* mutations observed (red) in the DDD data set versus the ones not observed (blue).

**Table 2:**
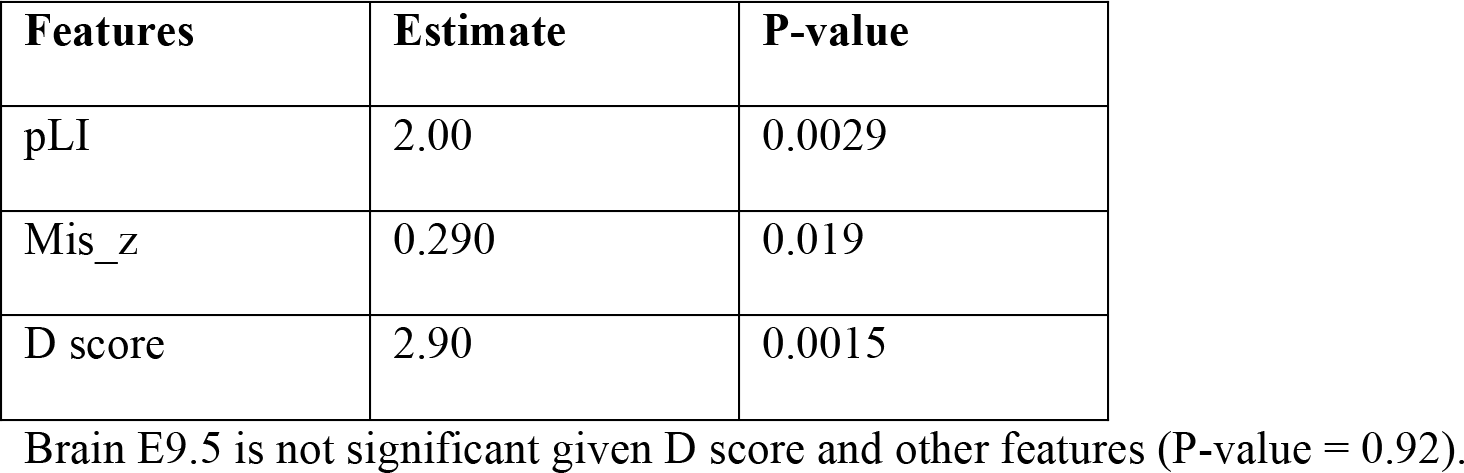
Logistic regression for classification of recurrently LGD-mutated genes in ASD and genes LGD-mutated in unaffected siblings.

| **Features** | **Estimate** | **P-value** |
| --- | --- | --- |
| pLI | 2.00 | 0.0029 |
| Mis_z | 0.290 | 0.019 |
| D score | 2.90 | 0.0015 |

Brain E9.5 is not significant given D score and other features (P-value = 0.92).

### Gender bias of autism-associated *de novo* mutations

The incidence of autism has a strong gender bias, with a male:female ratio around 4 overall and even higher for high-functioning cases. This is reflected in the population of participants included in the exome-sequencing studies (M:F=6.4 in the SSC dataset; Table S8). However, a lower incidence of *de novo* mutations in males was previously observed (10, 16, 17). We examined the gender bias of *de novo* LGD mutations in different sets of genes, focusing on mutations identified from the Simon SSC dataset, for which the number of male and female patients is known (Table S8).

Consistent with previous observations, a lowest M:F ratio was observed from genes with recurrent LGD mutations and genes with singleton LGD mutations predicted by the ensemble model (M:F~0.5, after correction for the gender bias of the participants; P<0.02, Fisher’s exact test). A more moderate, but significant, gender bias was observed in genes with singleton mutations predicted by D score alone (M:F=0.62; P=0.02, Fisher’s exact test). No significant gender bias was observed among singleton mutations in genes showing a negative D score (M:F=0.89; P=0.65, Fisher’s exact test). These observations provided an independent line of evidence that D-score and the ensemble model can discriminate disease-susceptibility genes from non-disease genes.

## Discussion

Here we present the use of cell type-specific gene expression profiles to improve prediction of autism-susceptibility genes. The molecular signature uncovered by DAMAGES analysis has several implications. First, this study echoes recent findings on convergent molecular pathways underlying autism etiopathogenesis including, approximately, three modules: synaptic structure and function, transcriptional regulation and chromatin remodeling, and Wnt signaling (reviewed by ref. (26) (18)). Conclusions of these studies were drawn from analysis of co-occurrence in genetic phenotypes (24), protein-protein interactions (14), and gene co-expression networks reflecting developmental dynamics in different brain regions (23, 25). This work extended these previous efforts by demonstrating that a robust signature of autism-susceptibility genes can be defined by their expression patterns in a range of specific CNS cell types (32, 33). Importantly, this signature reflects genes involved in transcriptional and post-transcriptional regulation and haploinsufficiency caused by LGD mutations in these genes. The importance of transcription factors and chromatin regulators in autism is now well established (14, 26). In addition, the role of post-transcriptional regulation is in line with the observation that several monogenic autism risk loci, including FMRP and MeCP2, are important regulators in RNA metabolism (58), and that candidate autism-associated genes show significant overlap with target transcripts of several neuronal RNA-binding proteins including FMRP (16, 59) and RBFOX1 (A2BP1) (43, 60, 61). Indeed, among the 822 FMRP target genes represented in the microarray dataset(59), a vast majority (718 or 87%) have a positive D score.

Second, while gene expression in brain or different brain regions was previously used to filter (13) or predict (62) candidate autism-susceptibility genes and, more recently, to reveal related pathways (23, 25), these previous methods were not optimized to predict individual autism-susceptibility genes or to provide rigorous assessment of the signal-to-noise ratio of resulted predictions. A key feature of DAMAGES analysis is that it adopts a case-control design using candidate genes derived from genomic DNA screens, which are completely independent of expression data. This design allows rigorous assessment of the biological relevance and predictive power of the uncovered signature by controlling potential confounding factors such as non-uniform mutation rates in different groups of genes. Our results demonstrated that the cell type-specific expression signature greatly increased the specificity of predicting autism-susceptibility genes with minimal loss of true hits. This is reflected in the observation that DAMAGES analysis predicted 35 out of 40 genes with recurrent LGD mutations identified so far and all 5 non-syndromic candidate genes with the highest confidence in the SFARI autism gene database. Importantly, information provided by the expression signature is complementary to the scoring metrics based on analysis of mutation frequency in the general population, and improved performance was achieved by combining the two methods.

Lastly, the cell-type specific signature captures a strong positive association with multiple types of cortical neurons, cerebellar granule cells, and striatal medium spiny neurons. This observation implies haploinsufficiency of genes that are normally highly expressed in these cell types as a converging pathogenic mechanism in autism. The implication of cortical projection neurons (25) and cells in the granule layer of the cerebellum (63) has been noted recently. In basal ganglia, striatal medium spiny neurons are known as the primary cell type vulnerable in Huntington's disease (64). In the context of autism, it has been shown that depletion of *SHANK3*, a gene highly expressed in striatum and regarded as the cause of the autism-related Phelan-McDermid Syndrome, results in ASD-like features such as impaired social interaction in mouse models (65). *SHANK3* has a singleton LGD mutation detected in the current exome-sequencing studies, and ranks among the top 5% genes genome-wide by the D score (see Figure 4A). In the cerebellum, a reduction of Purkinje cells and granule cells has been found in postmortem autistic brains and in mouse models (66, 67). Interestingly, our analysis revealed that cerebellar granule cells show a strong positive loading with a magnitude similar to those observed from cortical neurons, while Purkinje cells in cerebellum show weak loadings. These data suggest an intriguing hypothesis that different molecular mechanisms might underlie the loss of Purkinje and granule cells, although this has to be tested in future work. Finally, glial cells, especially astrocytes, show a strong negative loading. However, this should probably not exclude the contribution of these cells to autism. Instead, an alternative interpretation is that these genes may confer risks through other mechanisms than haploinsufficiency, which is supported by the observation that a subset of immune genes and glia markers are overexpressed in autism brains(43).

In summary, this study suggests the potential of utilizing gene expression and regulation information in predicting pathogenic mutations in autism. While we focused on cell type-specific expression in this work to demonstrate the proof of principle, we anticipate that additional expression profiles and functional annotations of genes, which can be easily integrated using a machine learning framework, will further improve the performance. Prioritized gene lists from such analysis can facilitate further validation by targeted re-sequencing in large cohorts (28) or more mechanistic studies using model organisms. This application will be particularly useful as the list of mutations are expected to grow steadily as a result of continuing autism exome- and genome-wide sequencing projects (4, 26) (27).

## Material and Methods

### Data compilation

For the current DAMAGES analysis, we used microarray gene expression profiles in 24 mouse central nervous system (CNS) cell types isolated from six brain regions, as well as unselected RNAs in each of these regions. This dataset was previously generated using a translational profiling approach named TRAP (34). For each gene, we selected the probeset with the maximum median expression across all 30 samples as a representative, if multiple probesets exist. In total, 20,870 genes with Entrez gene IDs are represented in the dataset. Expression intensities for each gene were first log2- transformed, with a pseudocount of 8 added to the intensities on the original scale.

The initial list of genes with *de novo* mutations in ASD probands and unaffected siblings were collected from four whole exome-sequencing studies (13–16). A total of 162 genes have *de novo* LGD mutations either in the probands or siblings. We determined the mouse ortholog of each human gene using the HomoloGene database(http://www.ncbi.nlm.nih.gov/homologene), complemented by manual searches. Mouse orthologs were found for 158 genes with LGD mutations, including 123 genes with LGD mutations only in probands and 35 genes with LGD mutations in siblings. Among these, a total of 145 genes, including 112 genes with LGD mutations in probands and 33 genes with LGD mutations in siblings, were represented in the microarray data, and were used for the initial model building and analysis. We also used 1,479 genes with log2 expression intensities ≥ 6 in two or more cell types and a standard deviation ≥ 2 across all cell types for PCA analysis shown in Figure 3B.

For further validation, we compiled an expanded list of genes with LGD mutations in autism patients (cases) or in their unaffected siblings (controls) from more recent exome-sequencing studies of about 3,960 cases and 1911 controls (17, 18). In total, we obtained a list of 670 genes, including 40 genes with recurrent LGD mutations in patients, 468 genes with singleton LGD mutation in patients, and 173 genes with LGD mutations in controls (note we excluded *TTN*). For genes that are not represented on the microarray data, we assigned their D score to the median value of all represented genes when we built the ensemble model (described below).

*De novo* copy number variation (CNV) data in ASD probands and annotations of overlapping genes were obtained from (11). This list is composed of 219 CNVs, and was compiled by the original authors from several previous studies (7–9, 11, 12). Technically redundant mutations, due to inclusion of the same patient samples in multiple studies, have already been removed from the list, so that recurrence of CNVs observed in the list is genuine. We similarly identified the mouse orthlogs of these CNV genes, and those (1,571 genes total) represented on the microarrays.

SFARI autism genes were downloaded in July 2013 from http://gene.sfari.org (38).

### Principal component analysis

DAMAGES analysis adopts a supervised classification framework taking advantage of the case-control design in the recent mutation screens. For the current work, the analysis is composed of two major steps: dimension reduction of gene expression data by PCA and prediction of autism-susceptibility genes by a regularized regression method.

For PCA analysis, log2-transformed expression intensities for each gene were first standardized across the 30 cell types to obtain zero means and unit standard deviations. PCA was performed in R using the princomp package. In brief, normalized expression values of each sample (i.e., cell type) were further standardized across genes to give a data matrix denoted as *X* = (**x**_1_,**x**_2_,… **x***_n_*) with *n* columns (genes) and *p*=30 rows (cell types). PCA (or equivalently, singular value decomposition) decomposes this matrix into the following form:

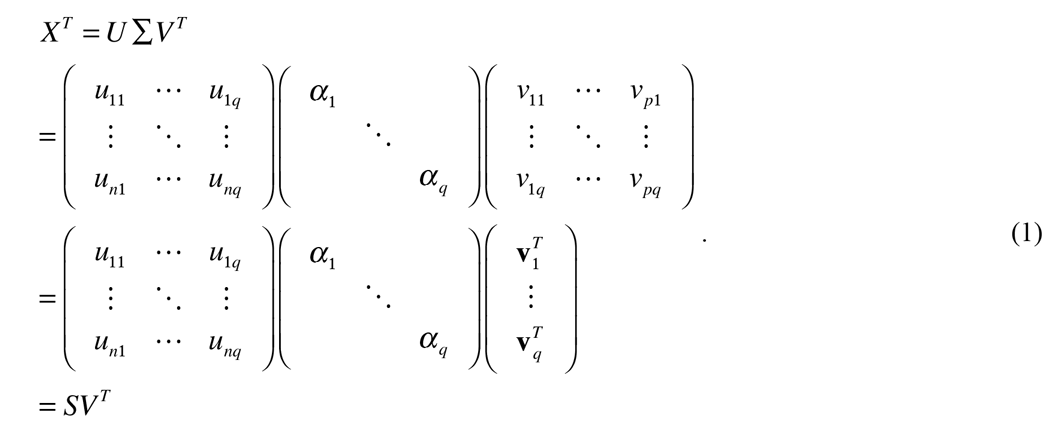

Here columns of *U*and *V* are unit orthogonal vectors (eigen vectors), and *α_l_ ≥ α_2_≥…α_q_* (q≥30). With this decomposition, each gene 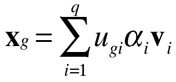, a linear combination of the first *q* principal components (PCs). Here *s_gi_ = u_gi_α_i_* is the score of gene *g* projected onto PC *i*; *v_ci_* is the loading of cell type *c* on PC *i*.

For a new set of genes, the score matrix is calculated by

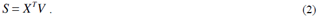

### Regularized regression analysis

We used a regularized regression analysis named lasso(37) to evaluate the contribution of each PC to prediction of each gene with respect to the source of mutations (probands versus siblings) with the following representation:

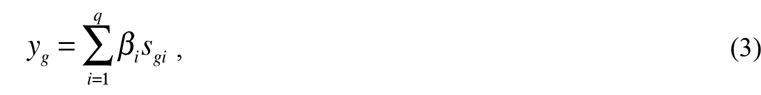

by minimizing

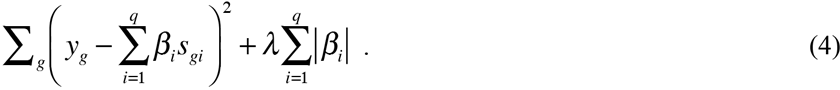

The second term controls the number of non-zero regression coefficients and therefore the model complexity. As the value of *λ* increases, the number of non-zero coefficients decreases. Model overfitting was evaluated by a standard leave-one-out cross validation (LOOCV) procedure, in which one gene was held out and the other genes were used to build the regression model and predict the gene left-out. For this study, we define the score *D_g_ = s_g2_* + 0.135 as the DAMAGES score or D score of gene g, in which the constant is determined by LOOCV.

### Estimating specificity and sensitivity in predicting autism-susceptibility genes

We first estimated the number of non-disease causing genes hit by random neutral mutations in the initial list of 112 genes with LGD mutations in probands (Table S1). The five genes with recurrent LGD mutations were considered to be highly confident true positives, given the very small chance to observe such recurrent *de novo* mutations (13, 16, 25). For the remaining genes, the number of non-disease causing genes, or false positives, was estimated based on the relative frequency of LGD mutations in siblings. A case-control design was used in three studies, and the number of false positives in each study was estimated separately. For one study (15), no sibling controls were included, so the number of false positives was estimated from the false discovery rate (FDR) of the other three studies pooled together. Overall, 53 genes with non-recurrent LGD mutations (42% of 127 genes) were estimated to be false positives. Therefore, we estimated that among the 112 genes with LGD mutations represented in the microarray dataset, there are ~65 disease-causing genes and ~47 non-disease genes, respectively.

To assess the single-LGD candidate gene prioritization performance of D score, ExAC metrics, and ensemble score, we used estimated background mutation rate (30, 68) to estimate precision and recall rate. Specifically, for each gene set (with G genes) defined by various metrics, we estimated the number of true positive (i.e. disease-causing; M_T_) LGD mutations based on the observed number (M_1_) of LGD variants in N cases and the expected number of variants (M_0_) given the background LGD mutation rate (*R_i_, i* indexes genes):

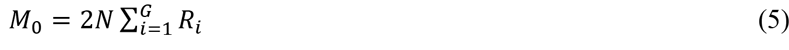

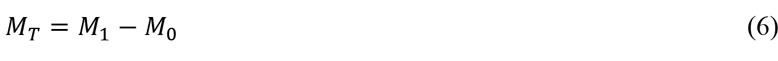

We denote the total number of true positives in all genes as M, and estimate sensitivity (recall) in each gene set by

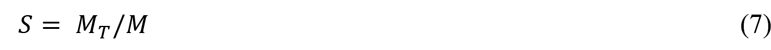

and precision by

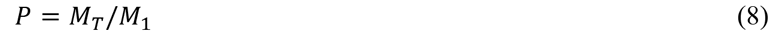

F-measure combines precision and recall by their harmonic mean:

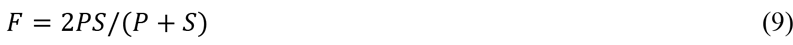

We note that genes with recurrent mutations in ASD patients and genes with LGD mutations in controls, which were used to build ensemble regression model, were not used to estimate the precision and recall in this analysis.

### Gene ontology and protein function analysis

Gene ontology (GO) analysis was performed using DAVID (41), using all protein-coding genes represented on the microarray as background.

## Statistical analysis

All statistical tests and logistic regression were performed using the R software.

## Acknowledgements

We would like to thank members of the Zhang and Shen labs for comments on the manuscript. This work was supported by grants from Simons Foundation (SFARI# 307711 to CZ) and NIH (R00GM95713 to CZ and U01HG008680 to YS). Computation was supported by NIH grants S10OD012351 and S10OD021764.

